# New Species of Anthurium (Araceae) from Huánuco Department, Peru

**DOI:** 10.1101/2025.01.31.635959

**Authors:** Armen Enikolopov, Thomas B. Croat

## Abstract

Two new species are described as new, fully characterized and compared with all other related congeners.

## INTRODUCTION

This paper is first in a series describing new species of Araceae from the Huánuco Department of Peru, with particular focus on the vicinity of Tingo María. Tingo María is located at the confluence of the Huallaga and Monzón rivers, where the eastern slopes of the Andes meet the western Amazon basin. Here, unique topography and orographic conditions create an isolated pocket of tropical wet forest, resulting in a functional biogeographic island.

At two locations in the Peruvian Andes, moisture-laden easterly air flow from the Amazon encountering the local topography creates patterns of orographic precipitation that result in high enough rainfall (to 5,000 mm/yr) to sustain areas of *Tropical Wet Forest* and *Premontane Wet Forest*, *sensu* Holdridge. The area immediately northeast of Tingo María is the northern-most of these. The next-nearest such biome occurs some 700 km to the southeast, in Madre de Dios, the second such location in Peru (Enikolopov, 2025) (Figure 1). Bounded by drier montane formations to the west and south and lowland Amazonian moist forest to the north and east, this effective insular isolation has likely contributed significantly to local speciation and endemism.

**Figure 1.**
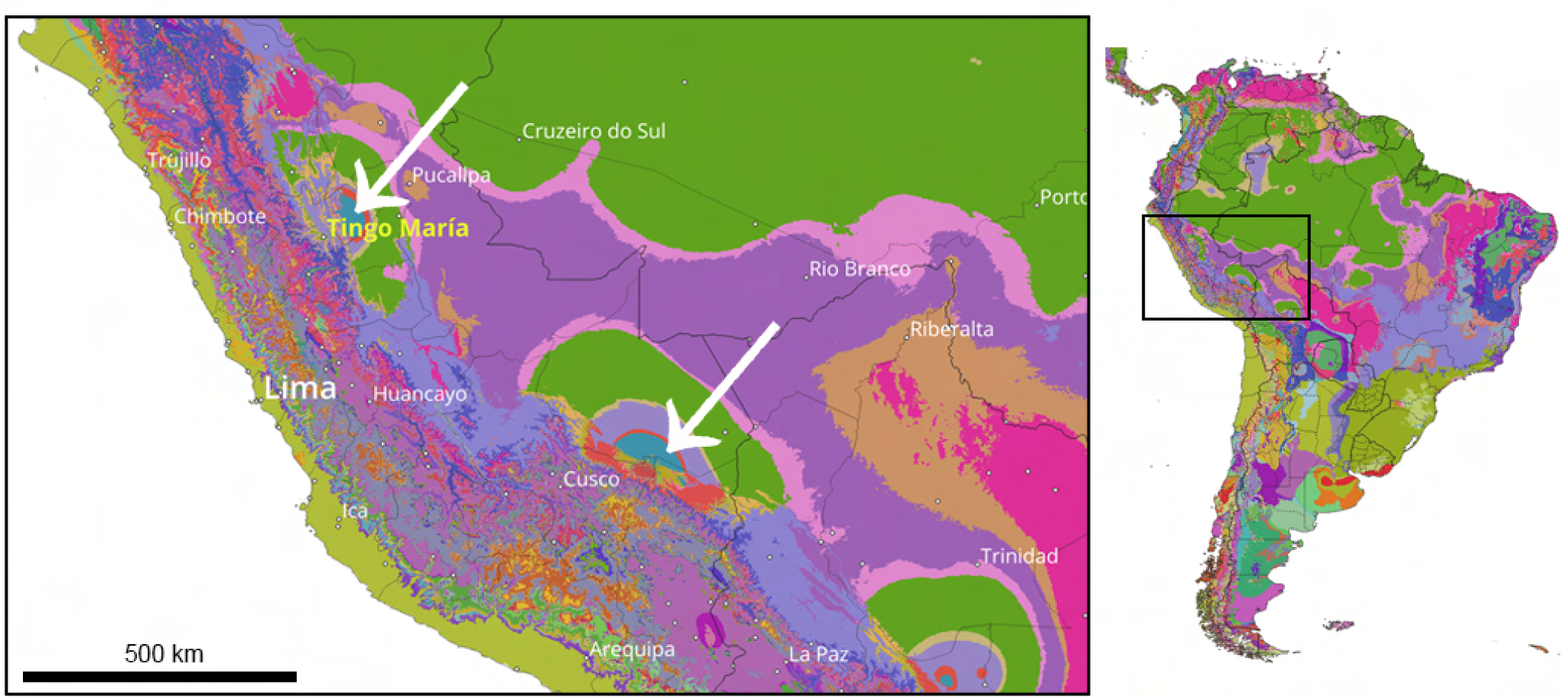
Functional insular biogeography. Holdridge Life Zone map, with transitional life zones, of South America with Peru inset, Tropical Wet Forest formations in cyan and marked with arrows. The northern (top) arrow marks the region near Tingo María, studied in this paper. The 700 km separation between these two isolated wet forest formations creates a biogeographic island at Tingo María. (Adapted from Enikolopov, 2025)

The Araceae of Peru are diverse, not primarily due to the total number of species, as it is exceeded in that category by Colombia, Ecuador, and Brazil, but rather due to the family’s diverse generic flora (Croat, 1999). In terms of generic diversity by country, it has the largest number of genera in the Neotropics. Owing to its geographic position, which straddles the tropical and temperate regimes, it has the largest number of ecological zones. In the case of the Holdridge life zone system, it is exceeded only by Bolivia in the number of life zones for a single country (Holdridge & al., 1971). Only the island of Borneo has more genera than Peru (Boyce & Croat, 2011).

The Araceae for the Flora of Peru (Macbride, J. Francis, 1936) included only 147 species in 12 genera (actually 13 since he included *Stenospermation* as a part of *Rhodospatha*). The Checklist for the Flora of Peru (Brako & Croat, 1993) published 57 years later treated 26 and 218 species, including two introduced genera, *Alocasia* and *Colocasia*. Considerable taxonomic work has been done since the publication on the Flora of Peru Checklist by the second author (Croat). These have included regional florulas such as that of the Iquitos area (Vásquez Martínez, 1997), the Flora of the Río Cenepa in northern Amazonas Department (Croat & al., 2005, 2010) and the Flora of Cerro Colán, in western Amazonas Department (Croat & al., 2021). Other works described new species of *Anthurium* (Lingan & Croat, 2005; Croat & Lingan, 2008; Croat & al., 2006; Martel & al., 2022), or the rediscovery of rare *Anthurium* (Croat & Lingan, 2005). Several papers described new species of *Philodendron* from Peru (Croat & al., 2012; Croat & Mines, 2022).

Botanical work on the Tingo María region spans nearly two centuries, beginning with Eduard Friedrich Poeppig (1798-1868), who arrived in Peru 1826, but intensive collection only became possible with the construction of road access in the 1930s. Despite 90 years of subsequent botanical work, recent surveys by the first author and long-term observations by Alfredo Loayza (pers. comm.) have shown that the Tingo María region remains rich in undescribed species of Araceae. Upcoming publications on new species of *Anthurium* and *Philodendron* by Croat and Fred Mueller, as well as work on *Anthurium* by Croat and Alfredo Loayza, expand our understanding of the region’s diversity, to which this paper adds two more species, *A. tenuilaminum* and *A. obtusicataphyllum*.

## MATERIALS AND METHODS

All descriptions presented here follow a pattern modified from that of (Croat & Bunting, 1979), with particular details ascribed to the surface features and morphology of the vegetation in addition to the characteristics of the inflorescence. A yet unpublished multichotomous Anthurium key, developed by the Royal Botanic Gardens Kew and the Missouri Botanical Garden using Lucid Builder (LucidCentral.org, Queensland, Australia), was used for species comparisons. Life zone ecology mentioned is based on the Holdridge life zone maps (Holdridge, 1967; Holdridge & al., 1971) and determined using the dataset and online tool recently finished by the first author (Enikolopov, 2025). Figure 1 was created with this same dataset and the QGIS (QGIS, 2024) software package.

### Conservation Status

The Red List status of both species described in this paper (I.C.C.N., 2024) is classified as Data Deficient (DD), as each is currently known from only a single specimen. This lack of information prevents an accurate assessment of their risk of extinction. Further research, particularly targeted field surveys and specimen collection, is necessary to understand their conservation status.

#### Anthurium tenuilaminum

Enikolopov & Croat, **sp. nov.** Type: Peru. Huánuco: Leoncio Prado; 32km NE of Tingo Maria on Ruta 5N to Pucallpa (19 km E of Puente Pumahuasi), in vicinity of the village of San Isidro, then 300 m S on dirt road to trailhead near water station. Small trail through wet forest, primary growth, steep slope, 09°13’18.70"S 075°49’46.30"W; elev: 1500 m, 24 July, 2024, *A. Enikolopov, A. Loayza & K. Bettelyoun 340* (holotype, MO-7102236 sheet 1 of 2!, MO-7102237 sheet 2 of 2!; isotypes, USM!, COL!, K!, B!). Figures 2–9

**Figure 2.**
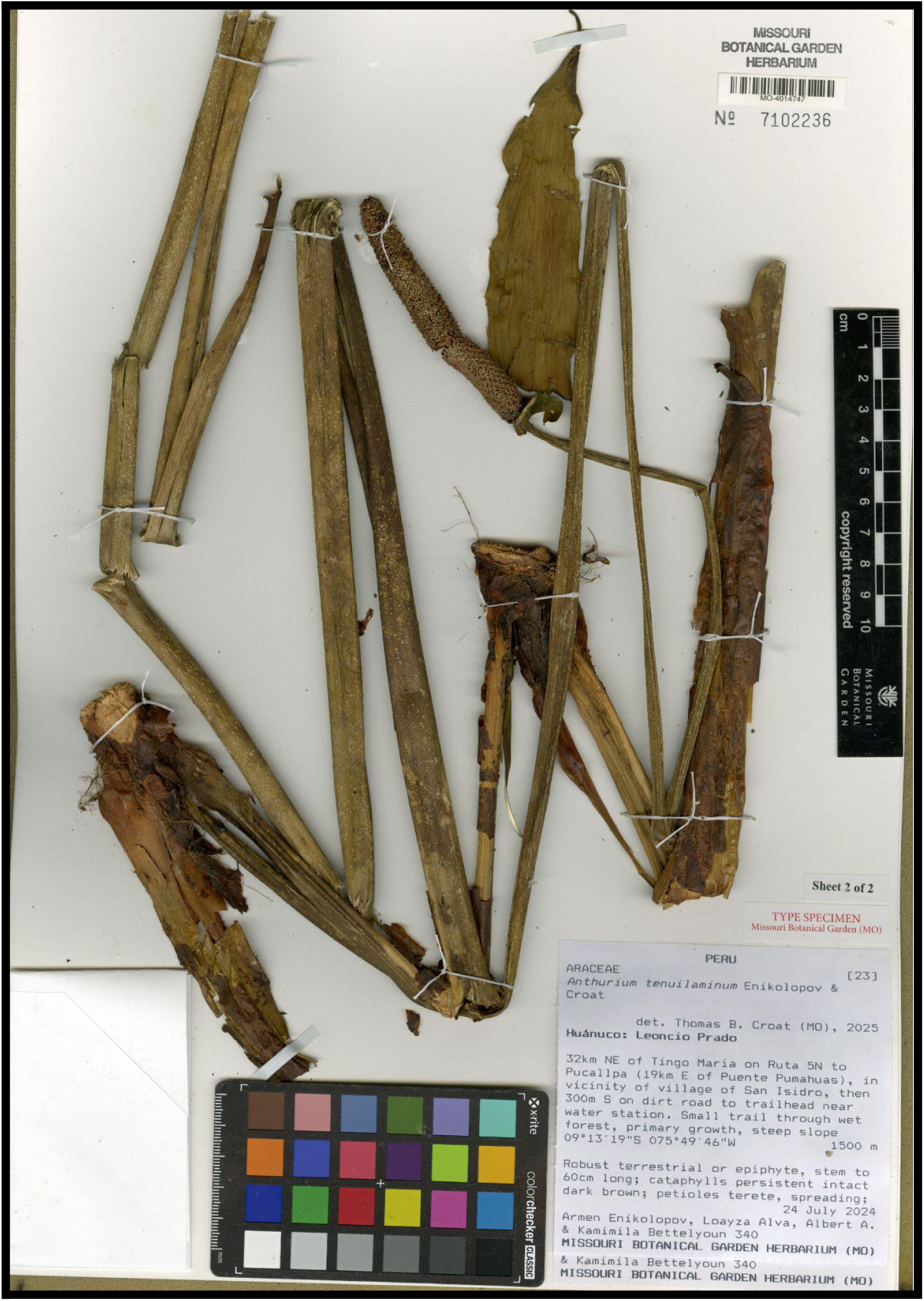
Anthurium tenuilaminum. Enikolopov & Croat, Holotype specimen: *A. Enikolopov, A. Loayza & K. Bettelyoun 340* (MO-7102236, sheet 1 of 2) showing stem, cataphylls, petiole and inflorescence.

**Figure 3.**
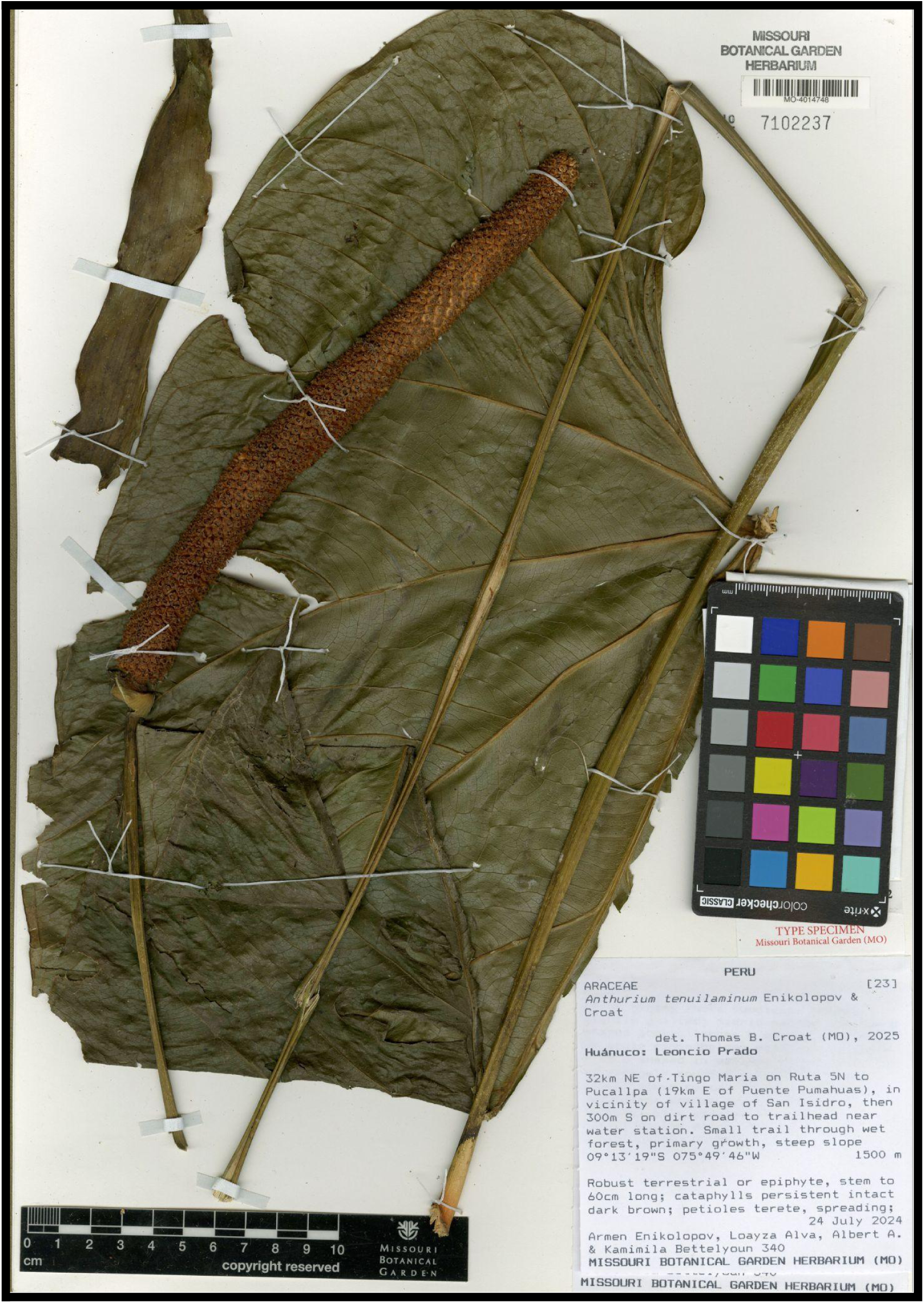
Anthurium tenuilaminum. Enikolopov & Croat, Holotype specimen: *A. Enikolopov, A. Loayza & K. Bettelyoun 340* (MO-7102237, sheet 2 of 2)

**Figure 4.**
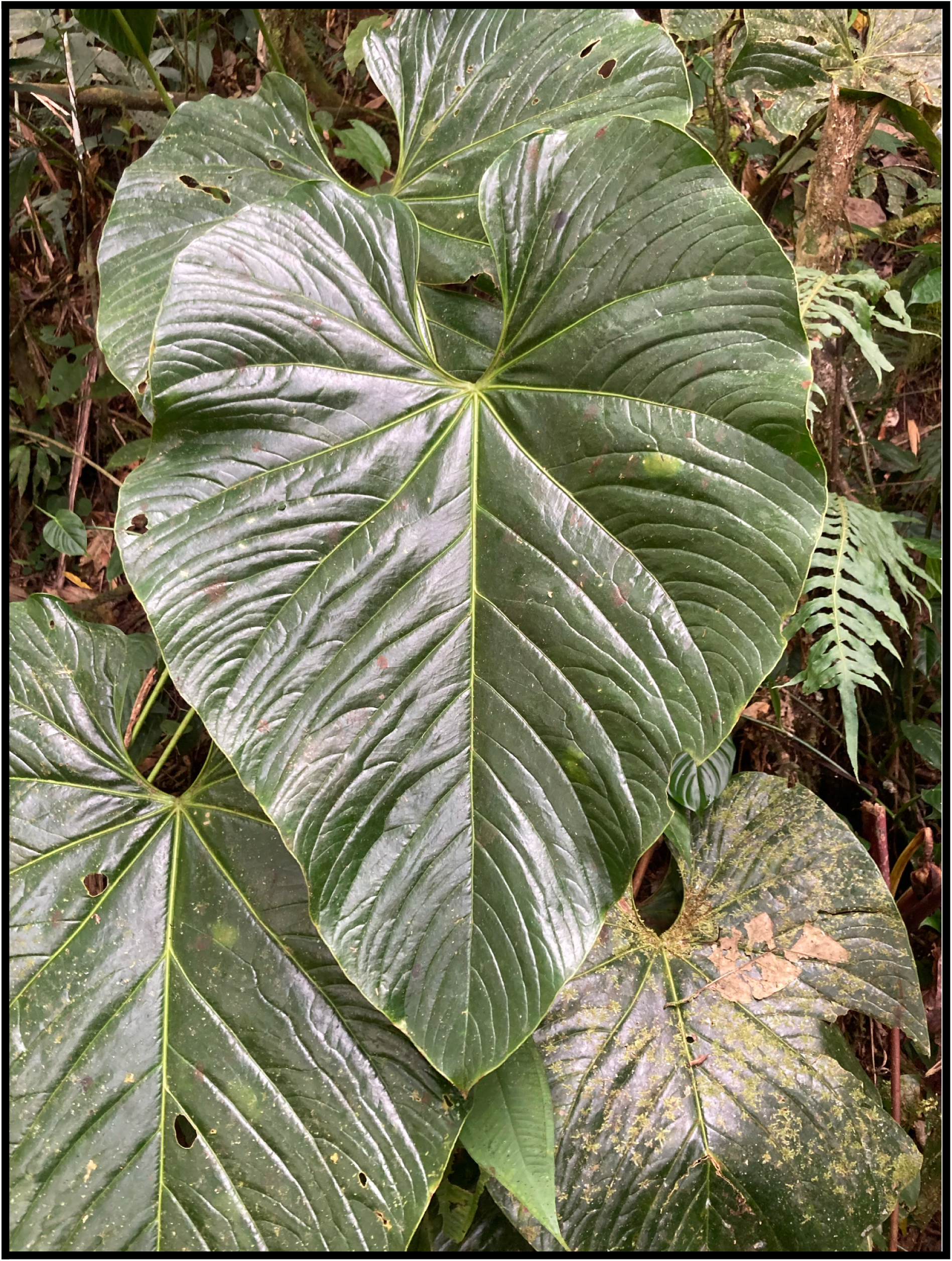
Anthurium tenuilaminum. Enikolopov & Croat, Adaxial blade surface, *in situ* at type locality.

**Figure 5.**
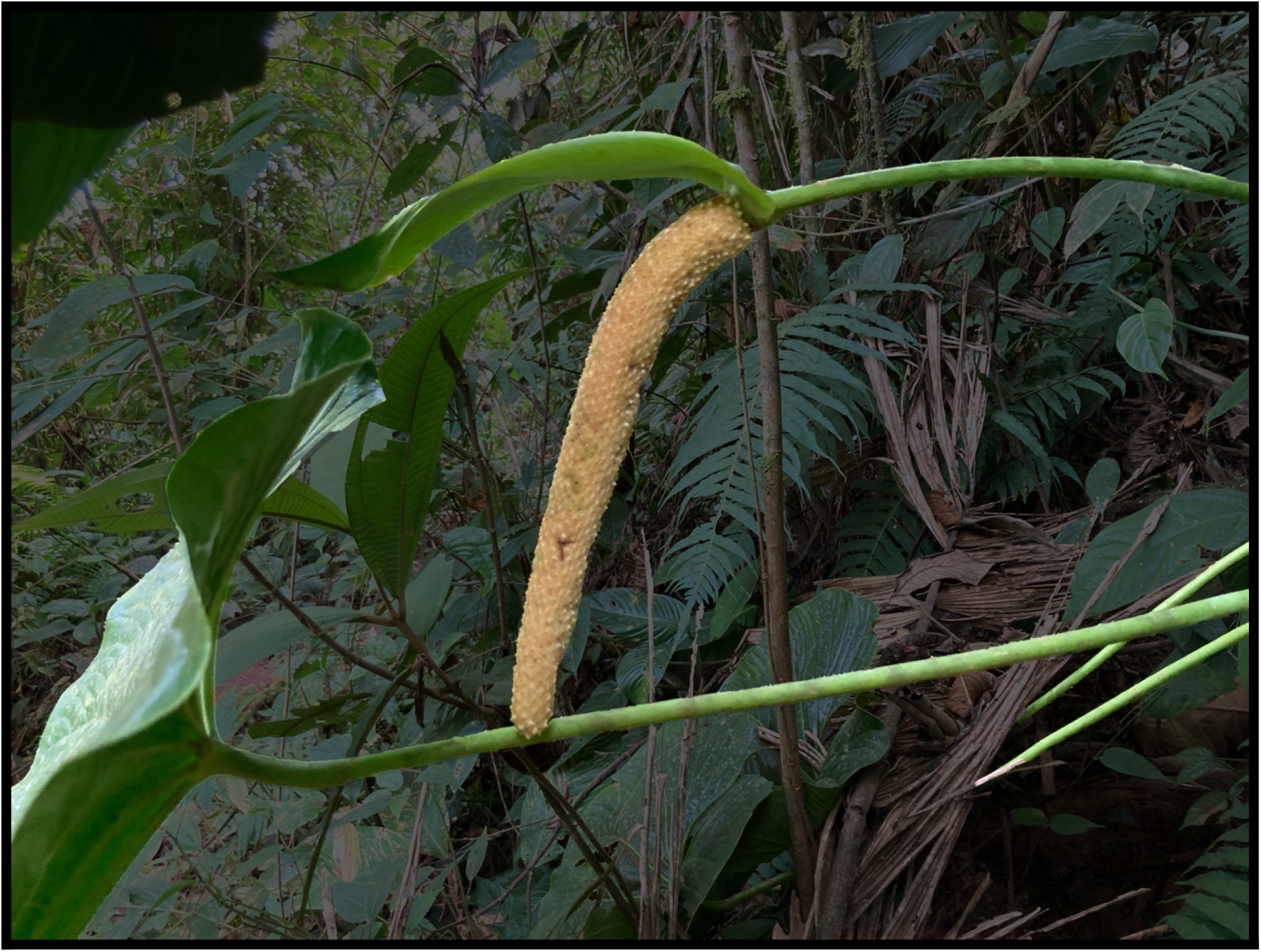
Anthurium tenuilaminum. Enikolopov & Croat, *in situ* at type locality. Note nodding spadix in early fruiting stage, hooded by the spathe.

**Figure 6.**
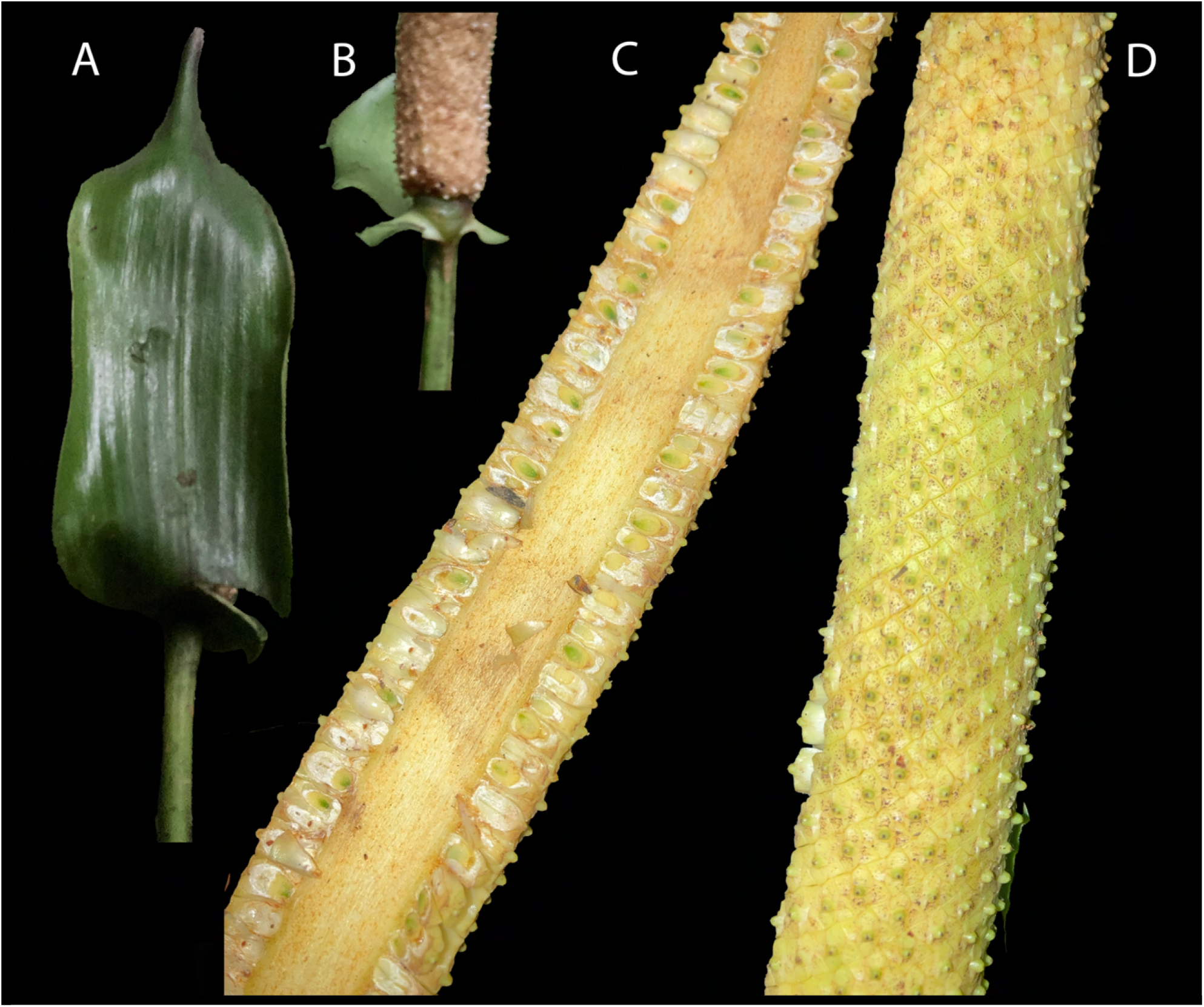
Anthurium tenuilaminum. Enikolopov & Croat, A. Spathe adaxial surface. B. Base of weakly stipitate spadix and front of spathe. C. Cut spadix of immature infructescence showing developing berries and weakly protruding styles. D. Outer surface of spadix of immature infructescence showing a few emerging berries. A & B to equal scale, C & D to equal scale.

**Figure 7.**
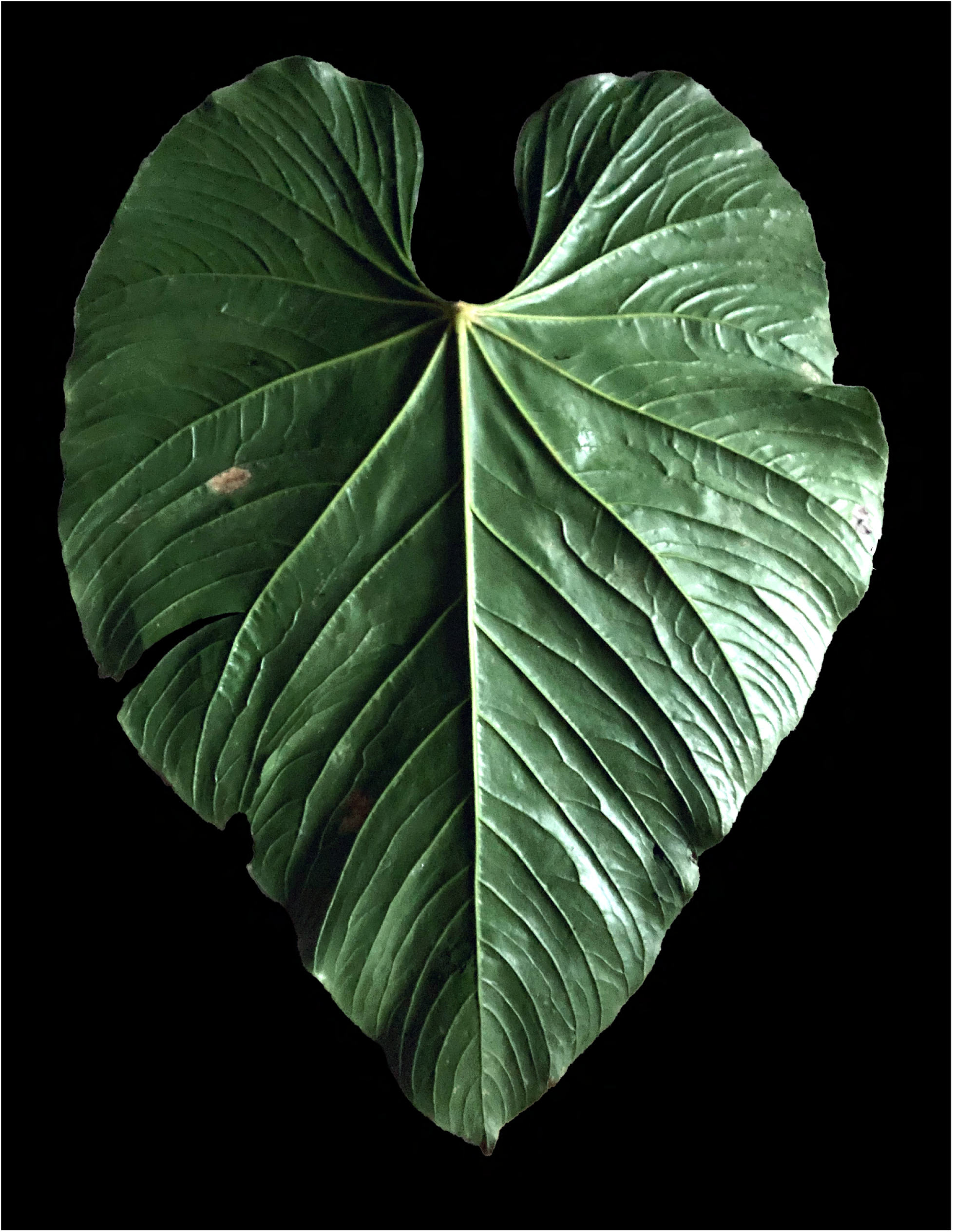
Anthurium tenuilaminum. Enikolopov & Croat, Abaxial blade surface.

**Figure 8.**
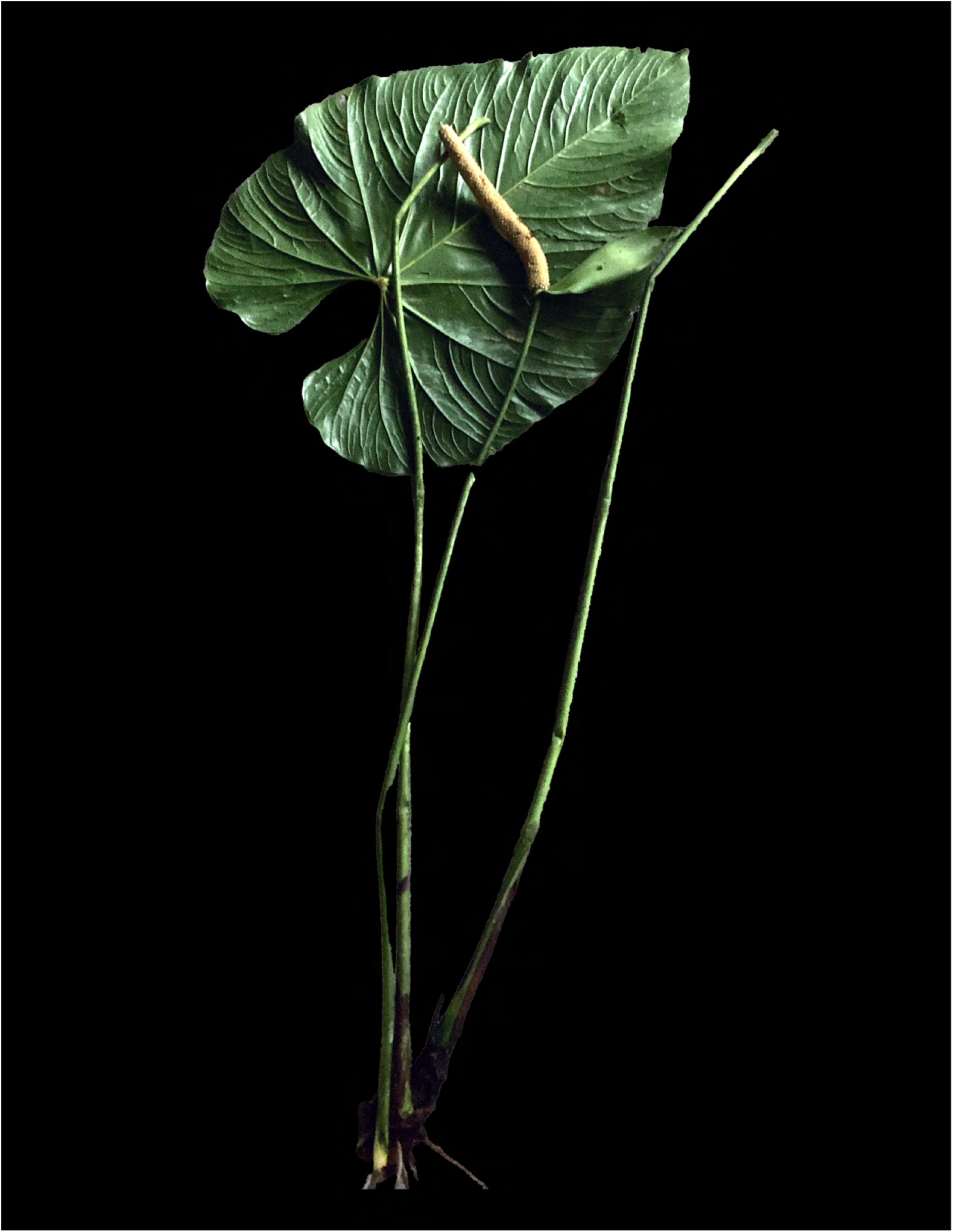
Anthurium tenuilaminum. Enikolopov & Croat, Leaf showing full petiole and leaf blade adaxial surface as well as a full inflorescence.

**Figure 9.**
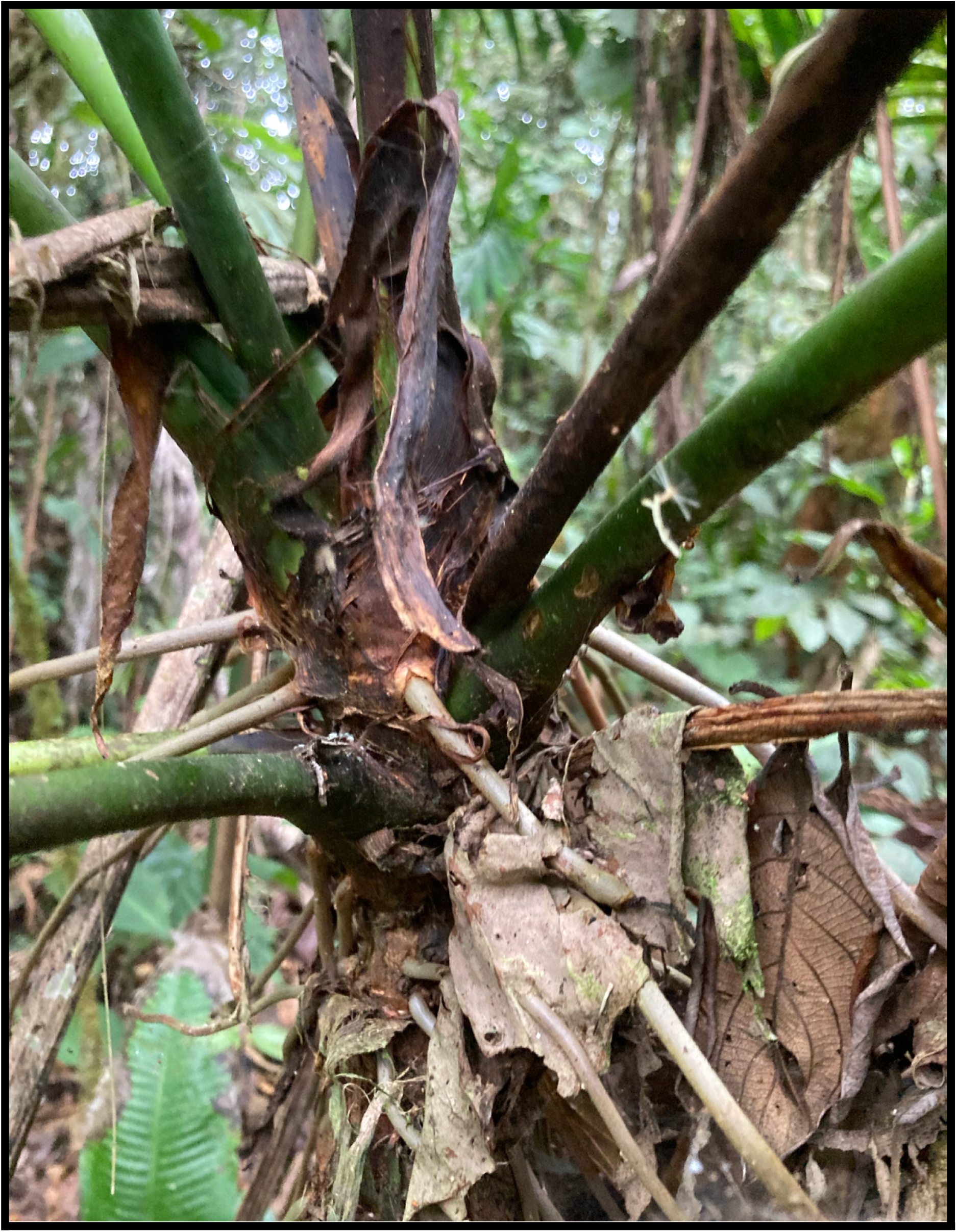
Anthurium tenuilaminum. Enikolopov & Croat, Apical portion of stem showing persistent intact cataphylls.

##### Diagnosis

The species a member of sect. *Calomystrium* characterized by its robust terrestrial or epiphytic habit, short internodes, terete non-sulcate petioles, thinly coriaceous large ovate-sagittate blades, narrowly hippocrepiform sinus, 7 pairs of basal veins, the 1^st^ & 2^nd^ free to the base, a nearly all naked posterior rib, 5–7 pairs of primary lateral veins,erect-spreading inflorescences, broad green erect-hooding spathe, and thick cylindroid curved, spreading or nodding, pale-yellow spadix.

Robust terrestrial or epiphyte, stem to 60 cm long; cataphylls to 19 cm long, persistent intact dark brown, drying moderately thin, dark brown, fragmenting at base with thin fibers; **petioles** terete, spreading, 82–86 cm long, drying to 1 cm wide midway, drying medium yellow-brown, weakly glossy, densely marked with short pale short-lineate excrescences (these filled with minute pale granular crystals); geniculum 2 cm long, slightly shrunken, darker, covered densely with rounded excresences as on shaft; **blade** 59–62 cm long, 40–45 cm wide, 1.3–1.5 times as long as wide, 0.49– 0.7 times as long as the petioles, thinly coriaceous, dark green and glossy above, weakly paler and subglossy below; midrib narrow shallow U shaped above, larger U shaped with rib or acute ridge below; upper surface diffuse-granular on magnification; lower surface weakly granular and sparsely gland-like-punctate on magnification; primary **lateral veins** 5–7 per side, arising at 35–40°, moderately quilted above; major veins above V-shaped to U-shaped in valleys above, U shaped or narrowly rounded below, drying bluntly acute and concolorous above, moderately acute and paler below; minor veins sunken above raised below, drying weakly prominulous below. INFLORESCENCE erect-spreading; peduncle spreading, 77–89 cm long, about as thick at the petiole; **spathe** green, oblong-lanceolate,19.5–21.5 cm long, 2.7–3 cm wide, spreading at 120° to the peduncle, then weakly arching, narrowly acuminate, hooding the spreading or weakly nodding spadix; **spadix** 8 cm long, 9–10 mm diam.; **flowers** 2.5–3 mm long, 2.2–2.5 mm wide; tepals gray crustiose-scaly; lateral tepals 1.2–1.3 mm wide, rounded on inside margin, 2–4(4-sided and shield-shaped); pistils weakly emergent, narrowly tapered and prominently exserted on drying, the stigma round, 0.1 mm diam.; INFRUCTESCENCE robust, 21.5–22 cm long, drying 1.7 cm diam. midway, cylindroid-tapered, nodding, pale yellow post-anthesis with traces of green; immature berries yellow with green ovules.

*Anthurium tenuilaminum* is endemic to Peru, known only from the type locality in Huánuco Department at 1500 m in a Tropical Premontane Moist Forest core life zone.

In the <u>Lucid Anthurium Key</u> the species tracks to *A. lutescens* Engl., differing by the absence of gland-like punctations on the lower laminar surface and petioles that are sulcate above and weakly keeled below; *A. cerratae* Croat & Lingan, differing by drying dark brown with a short, non-naked posterior rib and a long-tapered purplish stipitate spadix; *A. cerropascoense* Croat at 2000–2500 m from the Department of Junín which differs by drying dark brown, having only a single pair of free basal veins, a posterior rib only 3 cm long and naked 2 cm long as well as a green spadix; *A. consimile* Schott differing by its spathulate non-naked sinus and only 2 pair of primary lateral veins; *A. formosum* Schott, differing by its erect shrouding purple-tinged green spathe and its spindle-shaped purplish violet spadix with prominently exerted pistils and *A. rojasiae* Croat differing by its more coriaceous dark brown-drying blades with a collective vein much more distant from the margin.

##### Etymology

The species epithet comes from the Latin *tenuis* (thin) and *laminum* (blade) referring to the very thin-drying blades which is unusual for any member of sect. *Calomystrium*.

#### *Anthurium obtusicataphyllum* Croat, Enikolopov & Loayza, **sp. nov**

Type: Perú, Huánuco, Leoncio Prado, Along trail near stream through primary forest running within 5 m of stream. Trailhead starts across suspension footbridge 14 km S of Tingo Maria. 1.0 km from trailhead, 9°25’29.95"S, 75°58’36.12"W, 797 m, 27 July 2024, *A. Enikolopov, A. Loayaza & K. Bettelyoun 363* (holotype, MO-7102102!; isotypes, USM!, COL!, K!, B!). Figures 10–16

**Figure 10.**
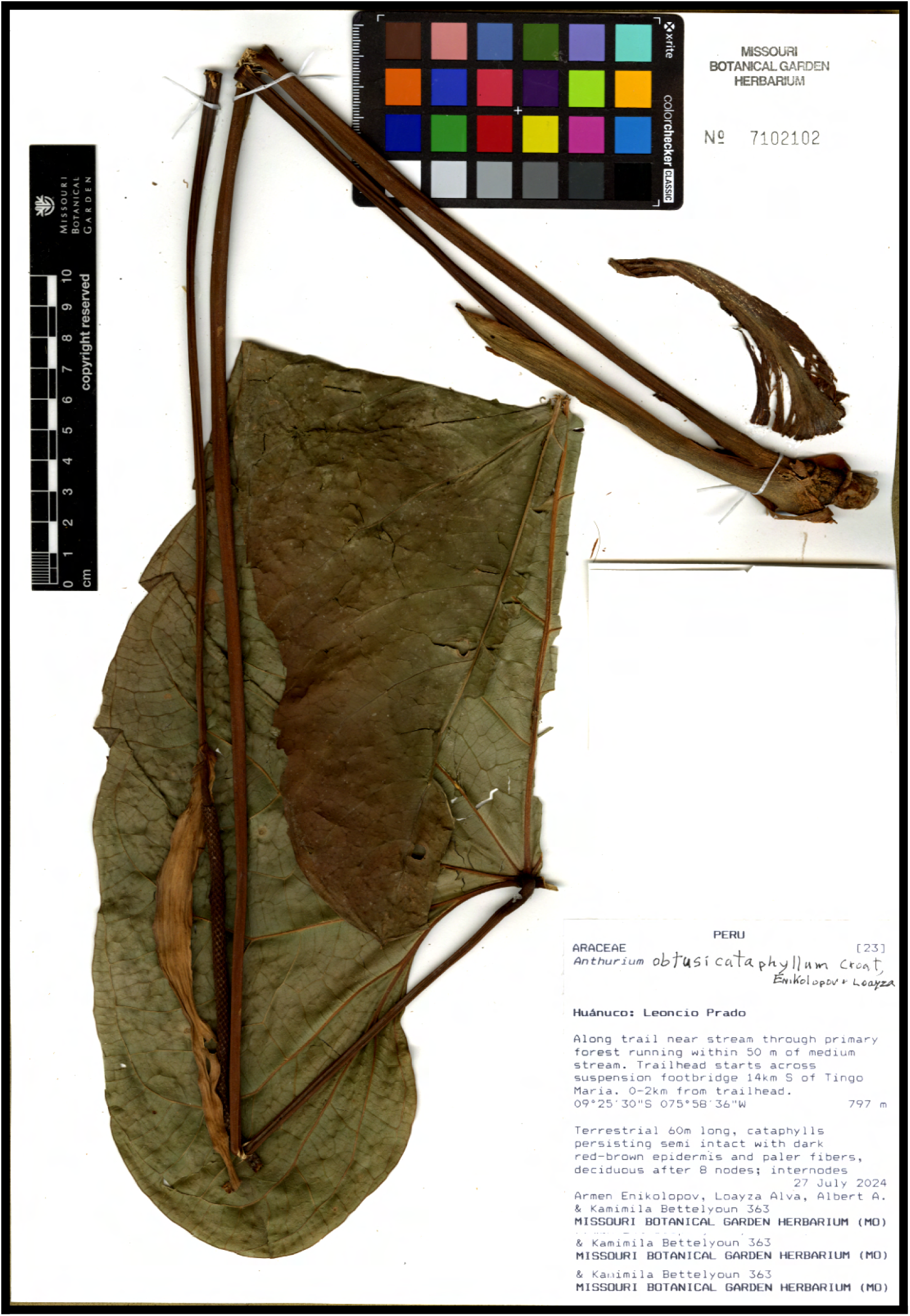
Anthurium obtusicataphyllum. Croat, Enikolopov & Loayza, holotype specimen (MO-7102102).

**Figure 11.**
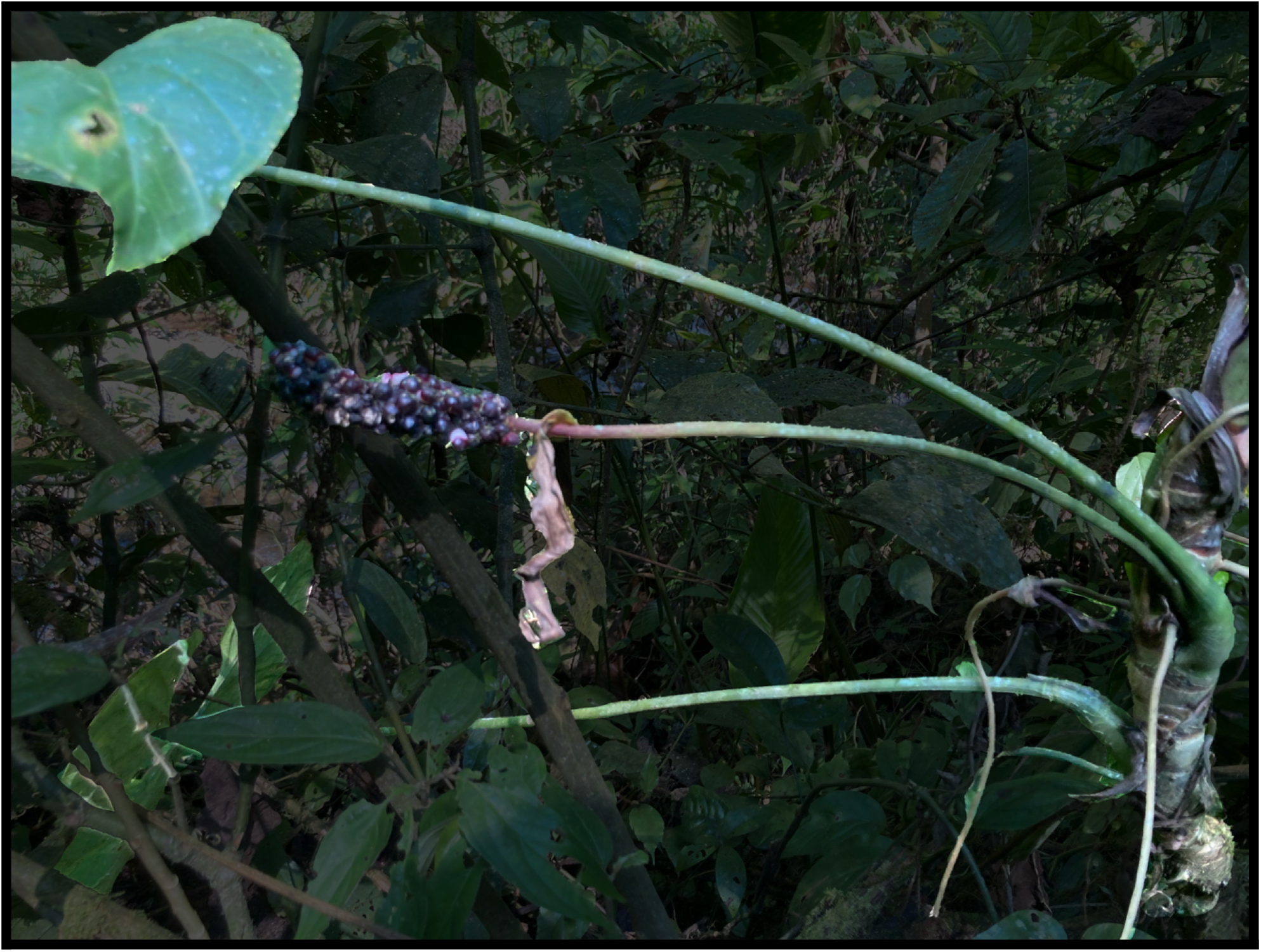
Anthurium obtusicataphyllum. Croat, Enikolopov & Loayza, habit, highlighting the spreading disposition of petiole and peduncle.

**Figure 12.**
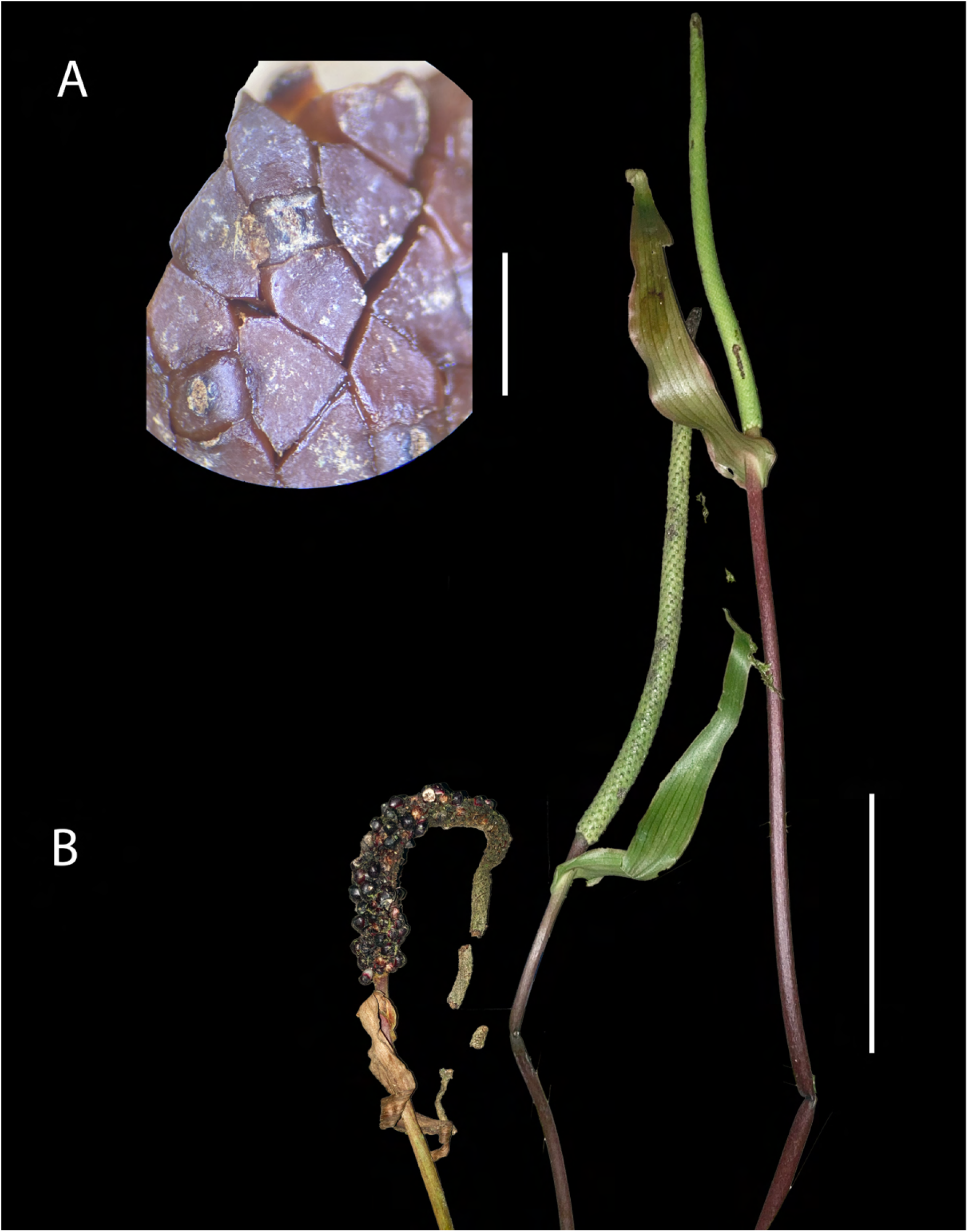
Anthurium obtusicataphyllum. Croat, Enikolopov & Loayza, A, Detail of alcohol-preserved immature infructescence. Scale bar = 2 mm. B. Infructescence (left), post-anthesis inflorescence (center), inflorescence at anthesis. Note violet peduncles. Scale bar = 10 cm.

**Figure 13.**
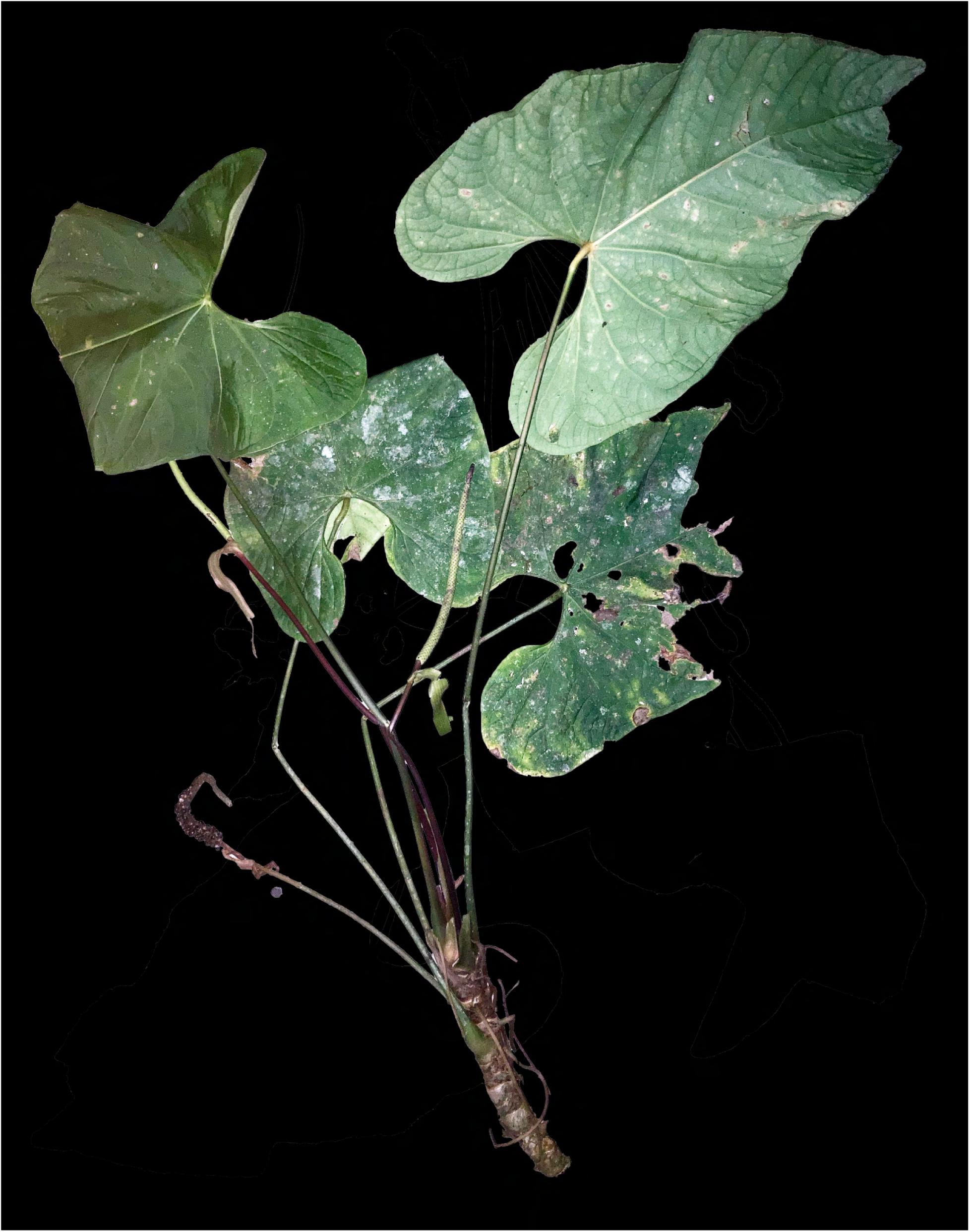
Anthurium obtusicataphyllum. Croat, Enikolopov & Loayza, Note violet peduncles.

**Figure 14.**
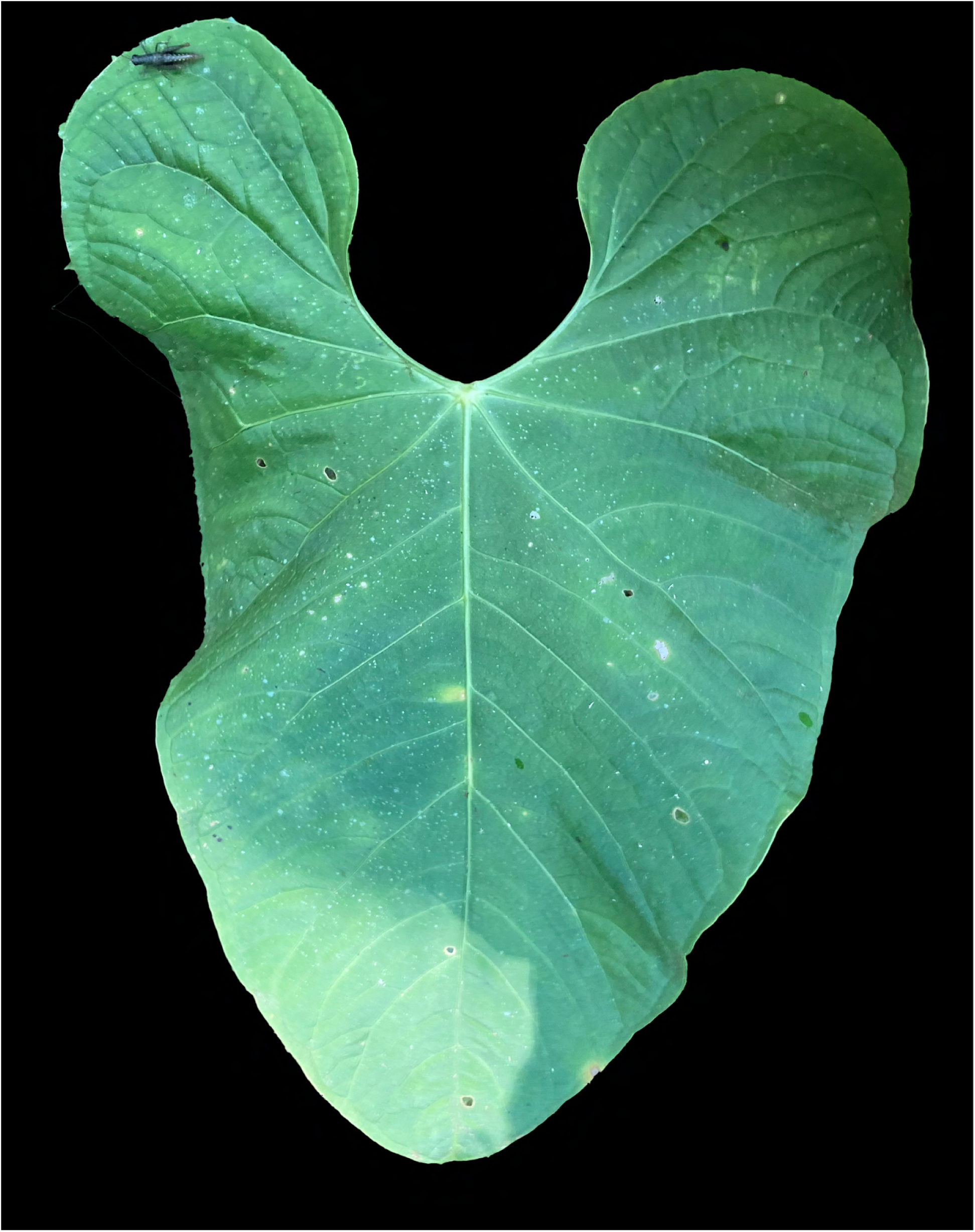
Anthurium obtusicataphyllum. Croat, Enikolopov & Loayza, Adaxial blade surface showing matte-subvelvet texture.

**Figure 15.**
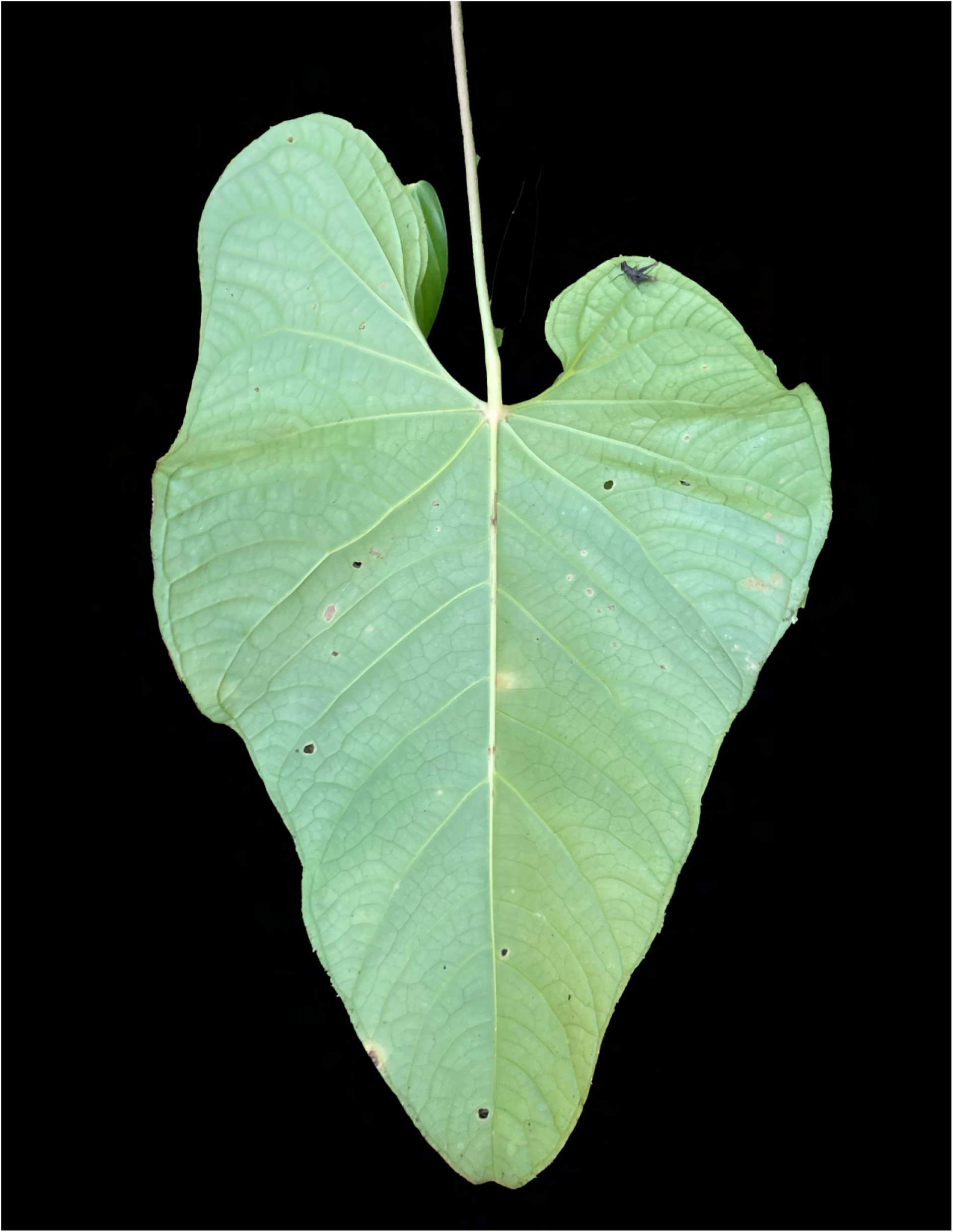
Anthurium obtusicataphyllum. Croat, Enikolopov & Loayza, Abaxial blade surface showing collective veins arising from primary lateral veins.

**Figure 16.**
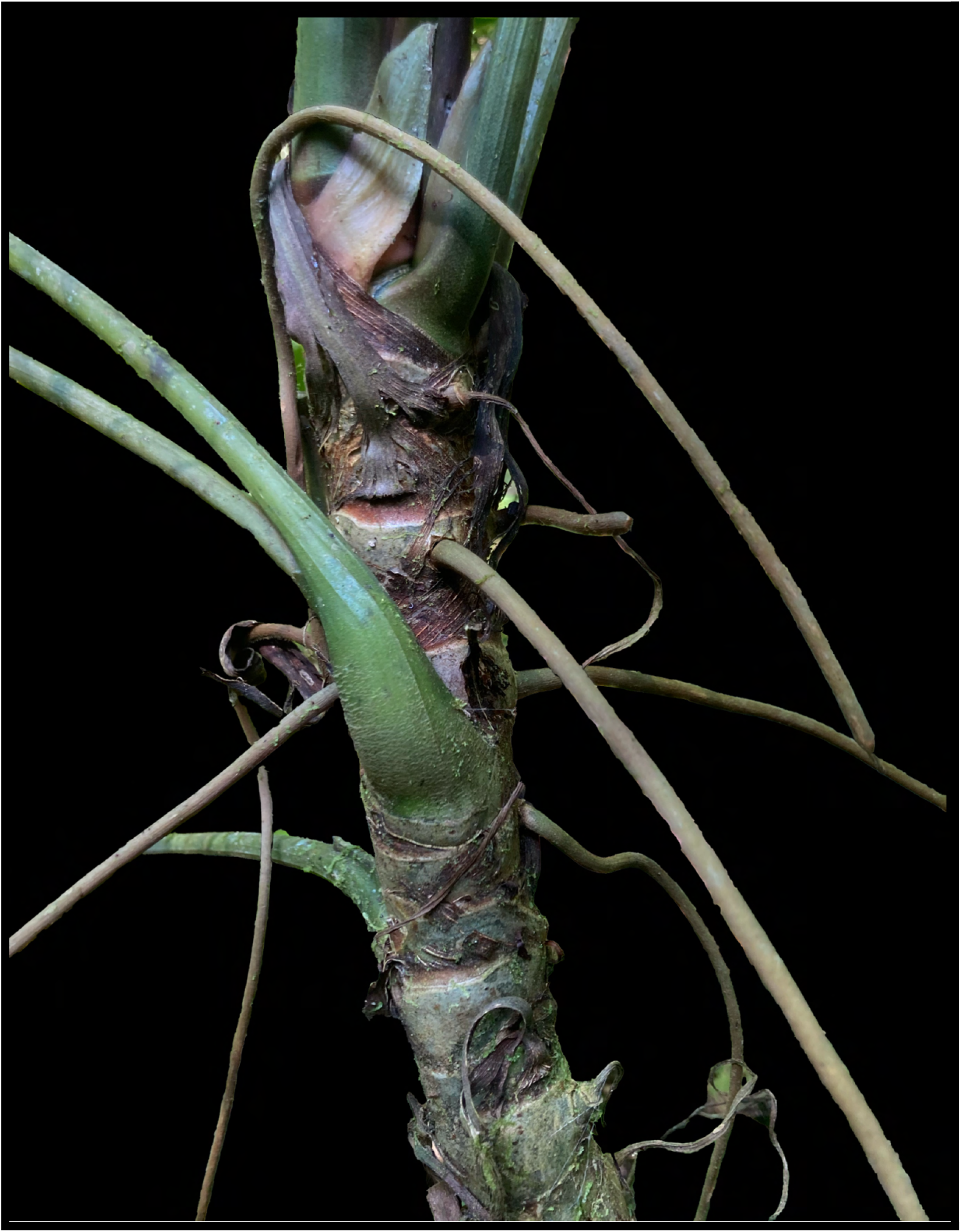
Anthurium obtusicataphyllum. Croat, Enikolopov & Loayza, Detail of stem. Note namesake blunt cataphyll at top of figure.

##### Diagnosis

The species is a member of *Anthurium* section *Cardiolonchium*, and is characterized by its terrestrial habit, semi-intact cataphylls, sulcate reddish brown-drying petioles, narrowly ovate-sagittate bluntly tipped matte-subvelvety blades 3-ribbed lower midrib, long-pedunculate inflorescence with a linear-lanceolate mostly green spathe and green long-tapered spadix with purple berries.

Robust terrestrial to 1.5 m; stem to 60 cm long, terete, gray-green, subglossy, with adventitious roots common at nodes, 3–4 mm thick (drying 0.9–1.6 mm), to 25 cm long, spreading just below horizontally, gray-green, matte, very weakly longitudinally striate in texture; internodes 2.5–3.5 cm long, diameter 2.5–3 cm, drying to to 2 cm, too; cataphylls 9–13.7 cm long, not ribbed, pointed and very narrowly rounded with both margins turned inward, with only a weak point at the apex less than 1 mm long, green sometimes tinged with shades of violet, persisting semi intact with dark red-brown epidermis and paler fibers visible basally, deciduous after ca. 8 nodes. LEAVES spirally arranged, ca. 5 in number, borne mostly at apical nodes; **petioles** erect to spreading, 48–79 cm long (average 66 cm), 6–8 mm diam., subterete, narrowly and shallowly sulcate adaxially with obtuse margins, sometimes with very multiple weak ribs, sometimes weakly transversely fissured at base, green to green with shades of red; drying reddish-brown finely striate, except basally, where pustular; sheath ca. 5 cm long, green with violet tone; geniculum 3–4 cm long, ca. 1/3 thicker than petiole, strong pale yellow; **blade** spreading-pendant on petiole, simple, ovate-cordate-sagittate, narrowly rounded and briefly short-pointed at apex, not at all acuminate, sagittately lobed at base, broadest near plexus, 42–50 cm long, 26–36 wide, 1.6–1.7x longer than broad, 0.75–0.83x as long as petiole; subcoriaceous and soft, matte-subvelvety above, submatte below, green above and below, drying moderately coriaceous, weakly bicolorous, medium greenish gray-brown, matte above, medium-light green, semiglossy below; anterior lobe 30–40 cm long, the margins convex, undulate; posterior lobes narrowly rounded, weakly inturned, 16–21 cm long; 10.3 cm wide midway; sinus hippocrepiform, 13–17 cm deep, 5–9 cm wide; midrib concolorous, U-shaped at base to ribbed at apex above, paler yellow-green, U-shaped and weakly 3-ribbed below, drying triangular, narrowly acute toward apex above, orangeish-brown below, drying round-raised, prominently 3-ribbed medium reddish brown below; primary lateral veins 4–8 pairs, acutely triangular above, narrowly raised, medially ridged below; departing midrib at 35–70°, curving towards apex; basal veins 7 pairs, the 1st acropetal, 2nd barely so; 3rd fused 2.4 cm, 4th 5th fused 4.7 cm, 6th & 7th fused 6.7 cm; posterior rib broadly curved back, 6–8 cm long, naked for 4–5 cm; tertiary veins weakly prominulous below; **collective veins** arising from 1st or 2nd primary lateral veins, weakly loop-connected, running 2–6 mm from margin. INFLORESCENCE erect to erect-spreading, shorter than leaves; peduncle 32–49 cm long (average 38 cm), 5–6 mm diam., 0.4–1.0 times as long as petiole, subterete, dark violet-red to maroon; spathe 15.5 cm long, 2–3 wide, linear-lanceolate, convex, smooth, soft, thinly coriaceous, green with red tinge and reddish-green margins, becoming medium green post-anthesis, subglossy, with ca 7. longitudinal veins darker than lamina, recurved, ca. ⅔ length of spadix, narrowly acute at apex, becoming weakly revolute at base post-anthesis, inserted at ca. 75°, margins meeting at base at a very obtuse angle; **spadix** gradually tapered, 17–22 cm long, diam. 8–19 mm at base, 4–5 mm at apex, light to medium green shortly pre- and post-anthesis, likely same at anthesis, stipitate, the stipe violet basally to green apically, 5–6 mm diam., 2.5 cm long in front, 0.8 cm long in rear; flowers sub-rhombic, sinusoid and parallel along secondary spiral, straight and parallel along primary spiral, 2.5 mm long, 2.8 mm wide post-anthesis, 7–8 flowers visible along principal spiral per side, 4–5 along secondary; lateral tepals 1.2–1.3 mm wide, inner margins rounded convex, outer margins 2-sided; stigma linear to ellipsoid, 1.3–1.4 mm wide, 0.45–0.55 mm long, 0.25–0.45 mm wide; stamens not observed INFRUCTESCENCE spreading, the apical portion dried and curled, lacking fruits; peduncle green to maroon; spathe persisting withered, reflexed, curled; spadix at least sometimes becoming recurved as berries mature; berries, dark near-black ruby, white at base, obovoid, to ca. 1 cm long, 7 mm diam.; seeds 2 per berry.

*Anthurium obtusicataphyllum* is endemic to Peru, known only from the type locality in Huánuco Department, Leoncio Prado Province, in the vicinity of Tingo María, immediately outside the southern border of Tingo María National Park at 797 m in a *Premontane Moist Forest* transitional life zone. The type was collected along a commonly-visited trail at an moderately sun-exposed location within 5 m of a stream.

The species is most easily confused with *A. sagittatum* (Sims) G.Don that differs by having a 5-sided to 5-winged petiole and the blades that are narrowly and gradually acuminate at apex (vs. narrowly rounded with short acute tip in *A. obtusicataphylum*. In the <u>Lucid Anthurium Key</u> the species tracks to *A. aylwardianum* Croat, which differs by its darker brown-drying blades with broadly spreading posterior lobes, a broadly parabolic sinus and a narrowly long-acuminate apex; *A. breviscapum* Kunth, differing by its much-elongated internodes, *A. oxapampense* Croat, differing by its panduriforme blades with sagittate-hastate lobes; *A. yungasense* Croat & Acebey from Bolivia, differing by its fibrous persistent cataphylls and its cylindroid spadix and A*. sanguineum* Engl. which differs by having a red spathe and dark green spadix.

##### Etymology

From the Latin *obtusus* (blunt) and *cataphyllum* (cataphyll), referring to the species’ blunt cataphylls.

## ACKNOWLEDGEMENTS

We thank Alfredo A. Loayza of Tingo María, Peru, for his assistance and knowledge of Peruvian *Anthurium*. We thank Kamimila Bettelyoun of New York for research assistance during field studies in Peru in the summer of 2024.

